# Meningioma microstructure assessed by diffusion MRI: an investigation of the source of mean diffusivity and fractional anisotropy by quantitative histology

**DOI:** 10.1101/2022.12.20.521068

**Authors:** Jan Brabec, Magda Friedjungová, Daniel Vašata, Elisabet Englund, Johan Bengzon, Linda Knutsson, Filip Szczepankiewicz, Pia C Sundgren, Markus Nilsson

**Affiliations:** Medical Radiation Physics, Clinical Sciences, Lund University, Lund, Sweden; Faculty of Information Technology, Czech Technical University in Prague, Prague, Czech Republic; Pathology, Clinical Sciences, Lund University, Lund, Sweden; Neurosurgery, Clinical Sciences, Lund University, Lund, Sweden; Russell H. Morgan Department of Radiology and Radiological Science, Johns Hopkins University School of Medicine, Baltimore, MD, United States; F. M. Kirby Research Center for Functional Brain Imaging, Kennedy Krieger Institute, Baltimore, Maryland, USA; Diagnostic Radiology, Clinical Sciences, Lund University, Lund, Sweden; Lund University Bioimaging Centre, Lund University, Lund, Sweden; Department of Medical Imaging and Physiology, Skåne University Hospital, Lund University, Lund, Sweden

**Author notes:** **Corresponding author** Jan Brabec, MD, MSc, PhD, Address: Barngatan 4, Skane University Hospital, 22185 Lund.

**Keywords:** Diffusion tensor imaging, Mean diffusivity, Fractional anisotropy, Cell density, Cellularity, Meningioma

## Abstract

**Background:** Mean diffusivity (MD) and fractional anisotropy (FA) obtained with diffusion MRI (dMRI) have been associated with cell density and tissue anisotropy across tumors, but it is unknown whether these associations persist at the microscopic level.

**Purpose:** To quantify the degree to which cell density (CD) and structure anisotropy (SA), as determined from histology, account for the intra-tumor variability of MD and FA in meningioma tumors. Furthermore, to clarify whether histological features other than cell density account for additional intra-tumor variability of MD.

**Materials and Methods:** We performed ex-vivo dMRI at 200 μm isotropic resolution and histological imaging on 16 excised meningioma tumor samples. Diffusion tensor imaging (DTI) was used to map MD and FA, as well as the in-plane FA (FA_IP_). Histology images were analyzed in terms of cell nuclei density and structure anisotropy (obtained from structure tensor analysis) and were used separately in a regression analysis to predict MD and FA_IP_, respectively. A convolutional neural network (CNN) was also trained to predict the dMRI maps from histology patches. The association between MRI and histology was analyzed in terms of coefficient of determination (R^2^). Regions showing unexplained variance (large residuals) were analyzed to identify features apart from cell density and structure anisotropy that could influence MD and FA_IP_.

**Results:** Cell density assessed by histology poorly explained intra-tumor variability at the mesoscopic level (200 μm) in MD (median R^2^ = 0.06, interquartile range 0.01 - 0.29) or FA_IP_ (median R^2^ = 0.19, 0.09 - 0.29). Samples with low R^2^ for FA_IP_ exhibited low variations throughout the samples and thus low explainable variability, however, this was not the case for MD. Across tumors, cell density and structure anisotropy were associated with MD (R^2^ = 0.58) and FA_IP_ (R^2^ = 0.82), respectively. In 37% of the samples (6 out of 16), cell density did not explain intra-tumor variability of MD when compared to the degree explained by the CNN. Tumor vascularization, psammoma bodies, microcysts, and tissue cohesivity were associated with bias in MD prediction when solely CD was considered. Our results support that FA_IP_ is high in the presence of elongated and aligned cell structures, but low otherwise.

**Conclusion:** Cell density and structure anisotropy account for variability in MD and FA_IP_ across tumors but cell density does not explain MD variations within the tumor, which means that low or high values of MD locally may not always reflect high or low tumor cell density. Features beyond cell density need to be considered when interpreting MD.

**Highlights:**

1. Cell density accounts for MD variability across but not within meningioma tumors.
2. Structure anisotropy accounts for in-plane FA variability across and within tumors
3. Vascularization, psammoma bodies, and microcysts influence the MD.
4. High and low meningioma tumor cell density can yield similar MD.
5. Features beyond cell density need to be considered when interpreting MD.

## Introduction

Diffusion MRI (dMRI) is the primary modality for obtaining information on tumor microstructure non-invasively (Stejskal and Tanner, 1965;Brown et al., 2014). Diffusion tensor imaging (DTI) is widely applied in patients with intracranial tumors and yields two key parameters: the mean diffusivity (MD) and the fractional anisotropy (FA) (Basser et al., 1994). MD correlates negatively with cell density (CD) in a wide range of tumor types (Sugahara et al., 1999;Gauvain et al., 2001;Chen et al., 2013;LaViolette et al., 2014;Surov et al., 2017). Decreased MD is therefore often interpreted as indicative of viable tumor regions with high CD. Furthermore, the FA reflects the voxel-level diffusion anisotropy and is generally high in white matter due to its highly anisotropic tissue structure. Therefore FA can be used to identify tracts displaced, disrupted or infiltrated by a tumor (Price et al., 2004;Yen et al., 2009;Jütten et al., 2019).

Although established on the whole-tumor level, it is not clear to which degree the correlation between MD and cell density, or FA and tissue anisotropy, holds quantitatively on a mesoscopic level within individual tumors. There are reasons to believe that both MD and FA can be affected by microstructural features other than cell density and tissue anisotropy. In the case of MD, cellular features such as size (Szafer et al., 1995), size of their nucleus (Xu et al., 2009), or membrane permeability (Colvin et al., 2011) are known to have an impact. MD can also be impacted by larger-scale mesoscopic features such as the presence of necrosis (Patterson et al., 2008) or stromal architecture (Squillaci et al., 2004;Yoshikawa et al., 2008). The features of the stroma may be tissue inhomogeneity, presence of large interstitial spaces, trabecula, nests and tubular formations or other complexity of intercellular spaces and junctions. Note that MD did not correlate with cell density in renal tumors and breast tumors (Squillaci et al., 2004;Yoshikawa et al., 2008). Furthermore, FA is known to merely reflect macroscopic (voxel-level) anisotropy, which is lower than the microscopic diffusion anisotropy due to the presence of orientation dispersion (Pierpaoli et al., 1996;Szczepankiewicz et al., 2016). This has been shown to be important in meningiomas, which tend to have high microscopic anisotropy but high orientation dispersion and thus low voxel-level anisotropy (Szczepankiewicz et al., 2016;Nilsson et al., 2020). Thus the interpretation of FA in meningiomas as an indication of tissue anisotropy could be biased by the orientation dispersion of the tumor microstructure (Szczepankiewicz et al., 2015;Brabec et al., 2022). Consequently, it is crucial to understand what affects MD and FA at the mesoscopic level when interpreting local changes of these parameters. However, few studies have investigated the relation between tumor microstructure as seen by microscopy to what is measured by dMRI on a voxel-to-voxel basis.

Meningiomas are the most prevalent primary intracranial tumor (34% of all intracranial tumors) (Louis et al., 2021). It has been proposed that DTI can be used for preoperative meningioma classification and consistency estimation, but results have been contradictory (Pistolesi et al., 2002;Gurkanlar et al., 2005;Hsu et al., 2010;Santelli et al., 2010;Lin et al., 2018;Yao et al., 2018). For example, some studies have shown that firm tumors are associated with lower MD values (Yogi et al., 2014;Miyoshi et al., 2020) or with MD values similar to gray matter (Romani et al., 2014). Other studies were not able to reproduce this result (Watanabe et al., 2016) or found that lower MD values are associated with variable consistency (Brabec et al., 2022). Furthermore, higher FA values have been associated with firm consistency (Kashimura et al., 2007;Tropine et al., 2007;Romani et al., 2014), suggesting that firm tumors may contain mainly anisotropic tissue with high microscopic diffusion anisotropy (Kashimura et al., 2007). Other studies, however, did not find such an association (Ortega-Porcayo et al., 2015;Brabec et al., 2022). DTI has also been proposed for differentiation of atypical, fibroblastic, and other meningioma subtypes (Jolapara et al., 2010;Surov et al., 2015), but both MD and FA have been reported as being similar across a wide range of meningioma types and grades (Brabec et al., 2022). To understand these divergent results, and if possible advise on ways to explain differences between studies, a better understanding of the link between meningioma microstructure and diffusion MRI results is needed.

In this work, we investigated the association between information derived from histology with that obtained from diffusion microimaging of the same specimen across six different types of meningiomas. We examined quantitatively to which degree cell density (CD) and structure anisotropy (SA) can account for the local intra-tumor variability in MD and in-plane FA (FA_IP_), respectively, as observed with dMRI with a voxel-to-voxel coregistered histology. The FA_IP_ is defined similarly to the FA but it disregards the trough-plane anisotropy making comparisons with thin histological slices more straightforward. Similarly to diffusion tensor analysis, the SA reflects the anisotropy in an image and is obtained from structure tensor analysis, which is similar to diffusion tensor analysis, analysis except that the diffusion encodings are replaced by spatial derivatives (Budde and Frank, 2012). To investigate if there were features beyond CD and SA that could explain MD and FA, we also trained a convolutional neural network (CNN) to predict MD and FA_IP_ from the histology slides. In addition, we qualitatively investigated voxels associated with large prediction errors in order to identify key microstructure features of meningiomas that drive a large change in the dMRI parameters.

## Materials and Methods

### Patients

This study included 16 patients with radiologically diagnosed meningioma tumors scheduled for surgical treatment between 2016 and 2018 at Skåne University Hospital, Lund, Sweden. Inclusion criteria were age above 18 years, histologically confirmed meningioma and signed informed consent. The study was approved by the Swedish Ethical Review Authority, and all subjects gave their written informed consent to participate in accordance with the Declaration of Helsinki. Table 1 and 2 provides a summary of the histopathological evaluation.

**Table 1.** Overview of histopathological classification of meningiomas samples. In total 16 samples were collected. The microstructural assessment was done according to the WHO criteria of 2016 (Louis et al., 2016).

| **SAMPLE** | **TYPE** | **GRADE** |
| --- | --- | --- |
| 1 | Transitional | I |
| 2 | Chordoid | II |
| 3 | Microcystic/Angiomatous | I |
| 4 | Meningothelial | II |
| 5 | Transitional | I |
| 6 | Meningothelial | II |
| 7 | Transitional | I |
| 8 | Meningothelial | I |
| 9 | Fibroblastic | I |
| 10 | Clear-cell | II |
| 11 | Transitional | I |
| 12 | Fibroblastic | I |
| 13 | Transitional | I |
| 14 | Microcystic/Angiomatous | I |
| 15 | Meningothelial | II |
| 16 | Transitional | I |

**Table 2.** Overview of histopathological classification. In total 16 samples have been investigated from which 11 were of WHO grade I and 5 of grade II. 6 different meningioma types were included and the most common was a transitional type of grade WHO I. Microstructural assessment was done according to the WHO criteria of 2016 (Louis et al., 2016).

| TYPE | GRADE | # |
| --- | --- | --- |
| Transitional | I | 6 |
| Fibroblastic | I | 2 |
| Microcystic/Angiomatous | I | 2 |
| Meningothelial | I | 1 |
| Meningothelial | II | 3 |
| Chordoid | II | 1 |
| Clear-cell | II | 1 |

### MR imaging and processing

In total 16 tumor samples were obtained after neurosurgical excision and fixated in formaldehyde solution (4%). The tissue was cut into blocks of approximately 35×20×2 mm^3^ (Figure 1A and 2B) to fit a 3D printed mold (Figure 2A) and scanned at a Bruker 9.4 T BioSpec Avance III scanner. DTI (Basser et al., 1994) was performed using a 3D-EPI sequence with TR = 2.5 s, TE = 30 ms, slices = 41, averages = 10, resolution=200×200×200 μm^3^, and with b-values of 100, 1000 and 3000 s/mm^2^ applied in six directions.

**Figure 1.**
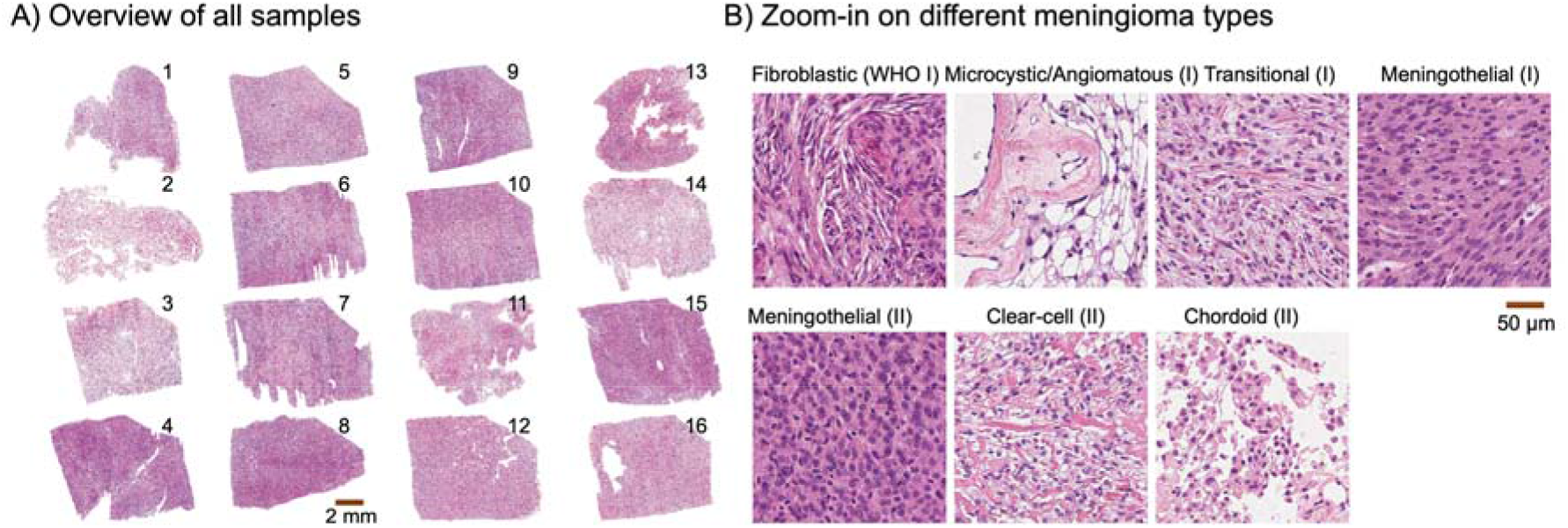
Histology overview. Panel A shows the 16 meningioma samples that were investigated. Panel B shows zoom-ins on different meningiomas types. Six were transitional, two fibroblastic, two microcystic/angiomatous, three meningothelial (WHO II), one meningothelial (WHO I), one clear-cell, and one chordoid. Microstructural assessment was performed according to the 2016 WHO criteria (Louis et al., 2016).

**Figure 2.**
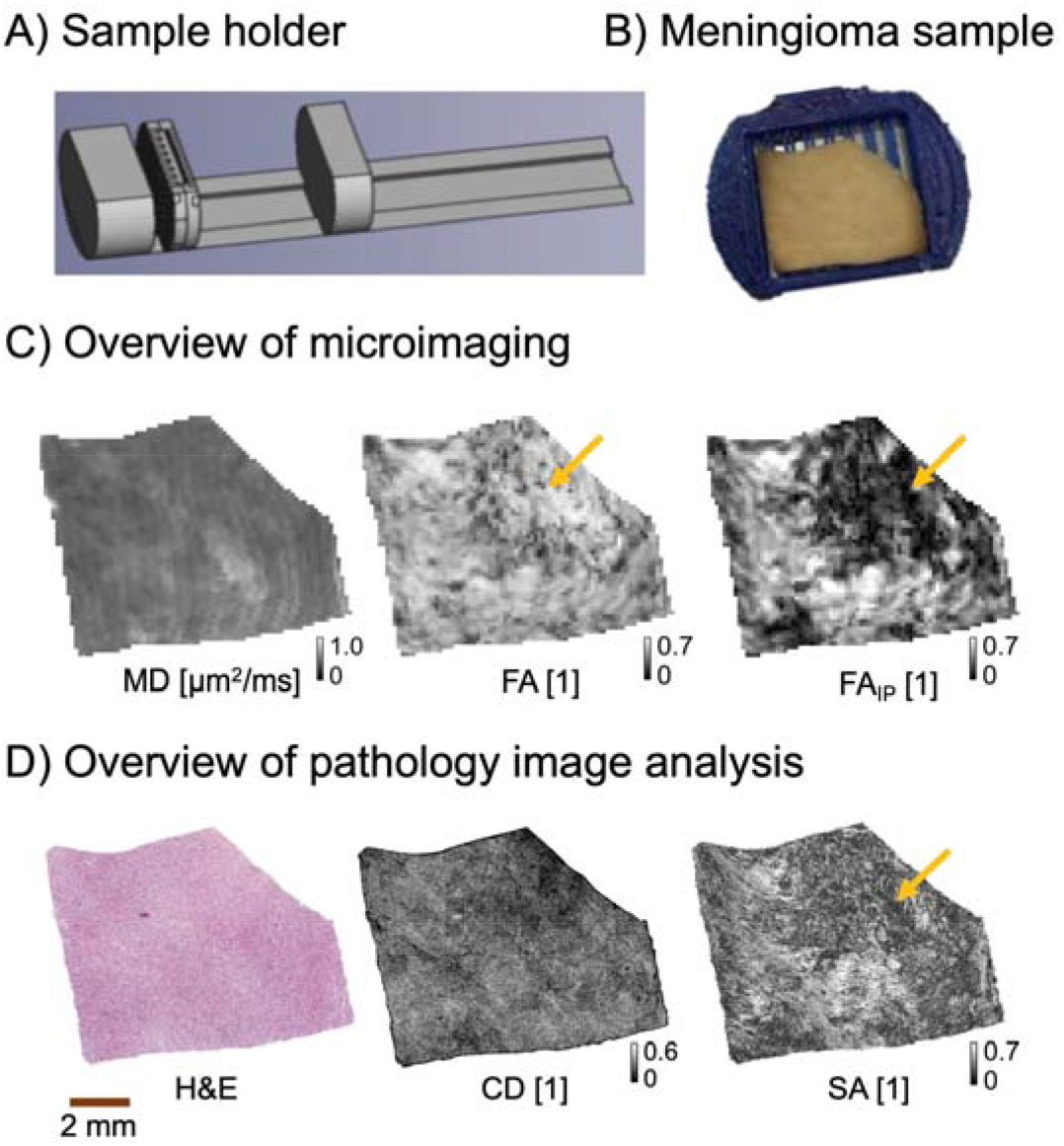
Methods overview. Panel A shows the schematics of the 3D-printed sample holder that was used to facilitate voxel-to-voxel coregistration. Panel B shows a meningioma (sample 5) in the holder. Panel C shows obtained dMRI maps: mean diffusivity (MD), fractional anisotropy (FA) and in-plane fractional anisotropy (FA_IP_). The latter captures only anisotropy within the imaging plane. The upper right part of the tumor has diffusion anisotropy that is dominant in the through-plane direction and therefore FA is high but FA_IP_ low (yellow arrows). Panel D shows a coregistered histology section (H&E stained) that was processed to obtain cell density (CD, cell nuclei count density) and structure anisotropy (SA from structure tensor analysis).

DTI analysis was performed with linear least squares fitting, as implemented in the multidimensional dMRI toolbox (Nilsson et al., 2018b) in order to extract maps of the FA, MD, and directionally encoded color (DEC) maps. Moreover, the FA_IP_ was calculated by utilizing only the in-plane eigenvalues of the diffusion tensor, according to

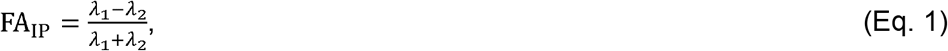

where *λ*_1_ and *λ*_2_ (*λ*_1_ > *λ*_2_) are eigenvalues of the diffusion tensor, reduced to in-plane (x-y plane) by setting *D*_xz_ = *D*_yz_ = *D*_zz_ = 0,

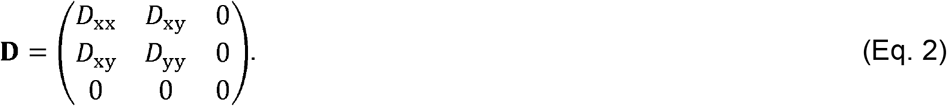

### Histopathology

The blocks on which MRI had been performed were embedded in paraffin, sectioned into 5 µm slices, and stained with hematoxylin & eosin (H&E). Each tumor specimen had been diagnosed for tumor type and malignancy grade. This diagnostic procedure adhered to the prevailing WHO criteria of 2016 as part of the clinical routine (Louis et al., 2016) because the data collection took place between the years 2016 and 2018. Sections were then digitalized at a resolution of 0.5×0.5 μm^2^. To facilitate coregistration, the sections were consistently taken from one side of the tumor block from the sample holder (Figure 2AB), which later allowed voxels from MR to be obtained from a similar location.

### Coregistration, cell density and structure anisotropy maps

H&E-stained histology images were coregistered to MR by, first, a rigid coregistration and, second, by a non-linear landmark-based approach. The landmarks were defined on the MD and FA_IP_ maps and then on the corresponding structures in the histology sections. Landmarks were placed at the corners and edges of the sections and also in tumor microscopic features, such as tumor microvasculature, readily discernible in both the histology sections and MR images.

Cell nuclei were segmented from H&E stained images using QuPath (version 0.23) cell detection algorithm (Bankhead et al., 2017). The segmentation was performed using the open-source code available at https://github.com/qupath. Furthermore, CD was obtained by exporting the cell nuclei centroid positions into a MATLAB environment where the cell nuclei counts were downsampled to match the MR resolution. This was achieved by summing of cell nuclei count over an area corresponding to a single MR voxel and, consequently, the CD map was normalized by dividing by the maximum CD value within the whole sample.

Structure anisotropy was obtained from a structure tensor analysis at the high histology resolution using on a previously described approach (Budde and Frank, 2012). This consists of computing a structure tensor **H** (Bigun, 1987;Budde and Frank, 2012),

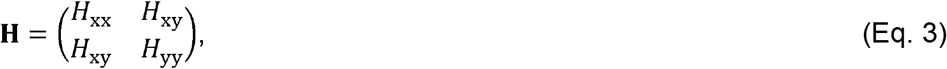

where *H*_xx_, *H*_yy_ and *H*_xy_ are partial spatial image derivatives along x or y directions. These were computed as convolutions of the histology image with derivative filters along either the x or y directions and blurred with a Gaussian filter (σ = 0.25 μm). Finally, the obtained structure tensor **H** was smoothed by another Gaussian filter (σ = 15 μm) and downsampled to match the MR resolution (200 µm) which was performed by averaging its eigenvalues within an area corresponding to a single MR voxel. SA was calculated from the eigenvalues λ_1_ and λ_2_ of the downsampled structure tensor **H** as

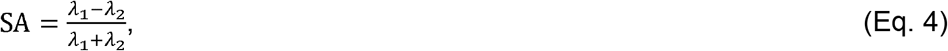

where λ_1_ > λ_2_. Finally, the SA maps were smoothed with the same Gaussian kernel as the dMRI maps (σ = 40 μm) to reduce the impact of small coregistration errors. Note that the calculation of SA and FA_IP_ are similar (compare Eq. 1-4).

### Prediction of MD and FA_IP_ from CD and IA

Since scatter plots between MD and CD or between FA_IP_ and SA values showed non-linear relationships (shown in Supplementary material Figure 1 and Figure 2), analyses were conducted to identify the function that best explained the relation between them. Five functions were tested: first-degree, second-degree, and third-degree polynomials, and in the case of MD, a second-degree polynomial constrained to be monotonically decreasing with maximal value at minimal CD to mimic the proposed negative association between CD and MD. In the case of FA_IP_, a first-degree polynomial constrained to the origin was tested instead. Results are shown in the Supplementary material Figures 1 and 2. It was observed that a second-degree polynomial was best suited in all cases and both modalities.

We quantitatively assessed to what degree CD and SA could explain the variability of the MD and FA_IP_, respectively, by using the coefficient of determination (R^2^) between the measured and predicted maps. This was calculated by randomly selecting 80% of the voxels as a training set. A second order polynomial in CD or SA was then fitted to MD or FA_IP_, respectively, for each modality and each sample. The explained variability in the remaining 20% of the voxels was then evaluated using R^2^. This procedure was used to unify the evaluation methods between the prediction by histology features and the one by the CNN (explained later). Furthermore, the process was repeated 1000 times with different random selections of the 80/20 split in order to estimate the uncertainty in R^2^.

We also investigated whether a lack of variability in MD or FA_IP_ within the sample could explain a poor association between intra-tumor predicted and measured dMRI maps. We quantified this by calculating R^2^ between R^2^ of the intra-tumor predicted versus measured MD or FA_IP_ and standard deviation of the MD or FA_IP_ across the sample, respectively.

### Quantitative comparison by convolutional neural network

We also quantified the variability in MD and FA_IP_ using a convolutional neural network (CNN) that was composed of an EfficientNetV2 network pretrained on the ImageNet dataset (Tan and Le, 2021) and fine-tuned with additional layers (network architecture overview in Supplementary material Figure 3). The CNN was designed to solve a patch-to-value regression task with the aim to predict either MD or FA_IP_ per voxel using a spatially corresponding patch of 360×360 color pixels from the histology images. We used horizontal and vertical image flipping for data augmentation and a train-validation-test split of 60/20/20%, 25 training epochs with early stopping and batch size of 32. The number of trainable parameters was 117 787 873. During training as a loss function the mean squared error was used and then the performance was evaluated on the test set using R^2^. The objective of this investigation was to determine whether a convolutional neural network could identify histological features that have an impact on the variability within MD or FA_IP_. Since the purpose was not to learn a general mapping from histology to DTI, the training and testing were conducted in a generous setting (within each sample rather than across all samples).

**Figure 3.**
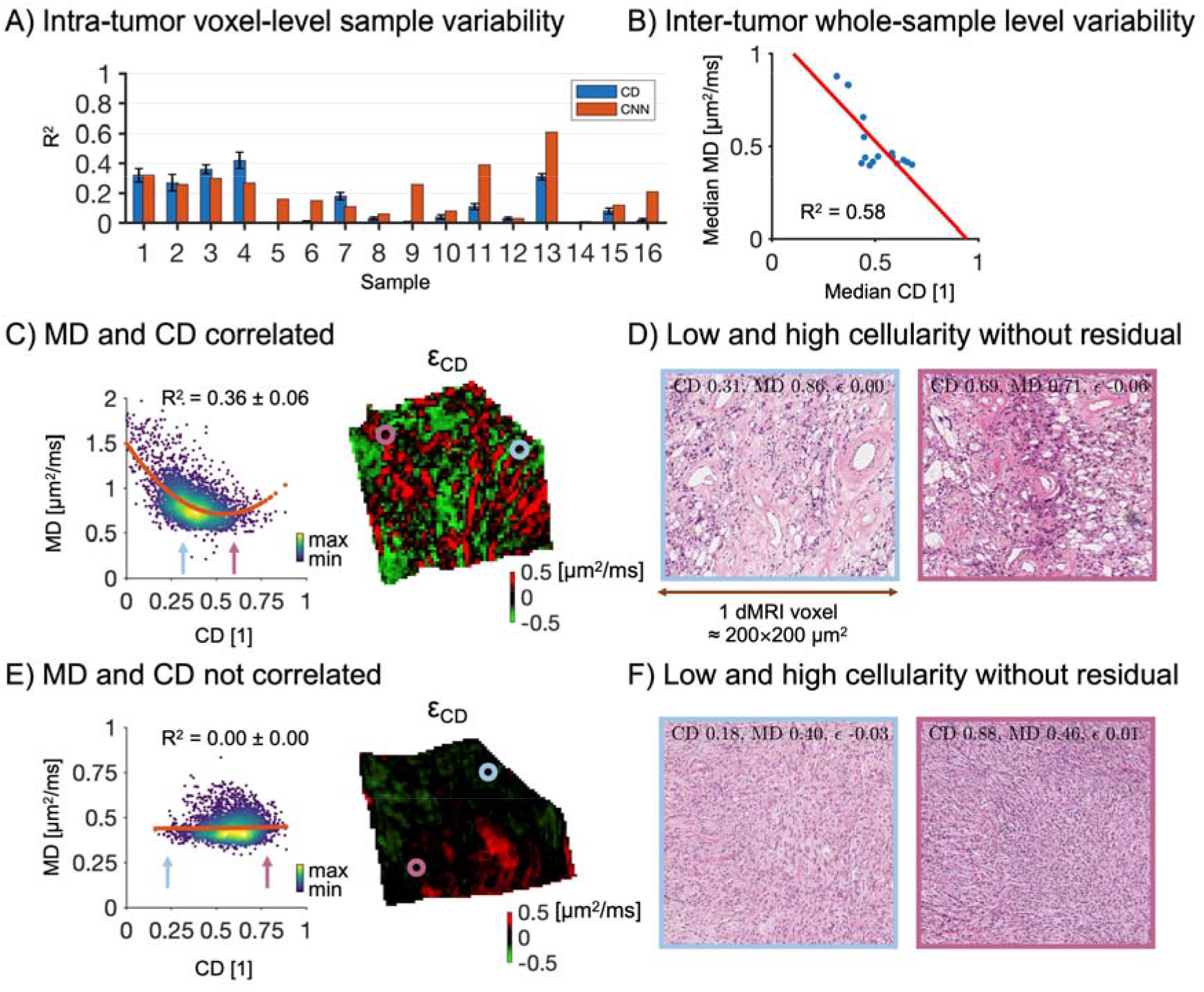
Association between MD and CD. Panel A shows the intra-tumor sample variability in MD explained by a second order polynomial in CD (R^2^; blue bars correspond to median, black error bars show interquartile range) and by the convolutional neural network (CNN; red bars), respectively. Panel B shows the intertumor whole-sample average of MD and CD (each blue dot corresponds to a single sample). The variability in MD across tumors is explained well by CD with R^2^ = 0.58. Panel C shows on the left a scatter plot from sample 3 where a strong correlation between MD and CD is present (R^2^ median ± interquartile range is displayed). High data density is marked by yellow color. A residual map is shown to the right. A voxel of intermediate CD (blue point) and another one with high CD (purple point) are indicated. Panel D shows their corresponding histology. The voxel with intermediate CD (blue inset) contains tumor stroma with vessels and microcysts, while the one with high CD (purple inset) has a clearer tumor mass and fewer microcysts and vessels. Panel E shows the same features but for sample 5. Here, the two voxels have similar MD despite having different CD (low versus high CD as indicated by the arrows in the MD-versus-CD plot). The voxel with low CD appears to have larger cells with larger cytoplasmic volumes than the cells in the voxel with high CD.

### Qualitative analysis by residual maps

To investigate additional features apart from CD contributing to the variability of MD, we studied residual maps (the difference between measured MD and predicted MD by CD). These maps were displayed using a color map where black corresponds to MR voxels without residual, green to voxels where the prediction was overestimated and red where it was underestimated. This approach revealed that the voxels where predicted MD was overestimated by CD (green color) would necessitate additional microstructure features causing “restrictions” with lower apparent diffusivity to counterbalance the effect of overestimation whereas the voxels where MD was underestimated (red color) would rather need a “free compartment” with high apparent diffusivity.

Similarly, we generated residual maps between MD predicted by the CNN and measured MD and compared them to those obtained by considering CD only. The purpose was to identify features impacting MD apart from CD. Attention was given to regions where the more general CNN approach had lower residuals than the less flexible CD-based regression approach, as this would indicate the CNN found in that region a feature to explain an MD deviation whereas the CD-based approach did not.

### Data and code accessibility

Analysis code and details of the MRI protocols are available at https://github.com/jan-brabec/microimaging_vs_histology_in_meningeomas. The dMRI data were processed by a software package for diffusion MRI available at https://github.com/markus-nilsson/md-dmri (Nilsson et al., 2018b). Additional data are available from the corresponding author upon request. Code for cell nuclei detection is available at https://github.com/qupath (Bankhead et al., 2017).

## Results

In total, 16 meningioma samples of six different types and two different grades were investigated (Table 1 and 2). An overview of sectioned blocks is shown in Figure 1, together with a display of the microstructural features of the different meningioma types. The histology and dMRI maps of MD, FA, and FA_IP_ were coregistered (Figure 2C). Note the difference between the conventional FA and FA_IP_ maps (Figure 2D). The latter captures the diffusion anisotropy within the imaging plane. Regions with high FA but low FA_IP_ indicate the presence of elongated cell structures pointing in the direction through the imaging plane. Examples of CD and SA maps obtained by analyzing histology slides (H&E stained) are shown in Figure 2D. Note that the SA map is highly similar to the FA_IP_ map but not to the FA map.

In the voxel-by-voxel within-sample analysis, CD poorly explained the intra-tumor variability in MD (Figure 3A), with R^2^ = 0.06 (0.01 - 0.29); median (interquartile range). The intra-tumor variability was better explained by the CNN, with R^2^ = 0.19 (0.09 - 0.29). In 37% of the samples (6 out of 16 samples; samples 5, 6, 9, 11, 13, and 16), CD explained much less of the variability than the CNN (the median ratio of the R^2^ of the CD vs CNN-based predictions was 7%). In the remaining samples, the CD-based approach explained a similar amount of variability as the CNN. The intratumor variability was weakly correlated with the standard deviation of MD within the sample (r = 0.51, p < 0.05, Pearson’s correlation coefficient), meaning that samples with lower variation in MD showed a weak tendency towards a weaker association with CD. When averaging the values of both CD and MD across the whole-sample and testing for an association across tumors we found a strong linear association with R^2^ = 0.58 (n = 16), although it is noteworthy that 5 out of the 16 samples with low CD and high MD stood out from the rest (Figure 3B).

To study the relation between meningioma microstructure and MD, individual samples were investigated. One sample showed a clear negative association between CD and MD within the sample (R^2^ = 0.36, Figure 3C). A closer inspection of two voxels with either intermediate or high CD is shown in Figure 3D. The voxel with intermediate CD and higher MD contains tumor stroma, vessels, and microcysts whereas the one with high CD and lower MD is characterized by a clearer tumor mass and fewer microcysts and vessels. Another sample showed no discernible association between MD and CD (R^2^ = 0.00, Figure 3E). A closer inspection of two voxels with similar MD but either low or high CD from that sample showed that the one with low CD contains cells with a larger cytoplasm volume than the one with high CD (Figure 3F).

To understand which features could affect MD beyond CD, residual maps were examined. This procedure led to the identification of five types of microstructure features of importance to MD. First, tumor vasculature was associated with an underestimated MD. This is shown in Figure 4A where an MRI voxel associated with a high residual shows the presence of vessels (histology with blue border) while a voxel with low residuals lacks them and rather features a solid tumor mass (purple border). Second, tightly packed microcysts were associated with an overestimated MD. This is shown in Figure 4B, where a voxel containing microcysts (blue border) is compared to a voxel with a denser tumor mass (purple border). This indicates that microcysts act as diffusion restrictions similar to cell bodies. Furthermore, the overestimation from CD is not present when MD is predicted by the CNN (Figure 4B, residual map □_CNN_), indicating that the CNN to some extent captures microcysts as a relevant feature. Third, psammoma bodies were associated with an MD overestimated from CD (Figure 4C). Similar to the case for the microcysts, this bias is absent for the prediction by CNN. Finally, tissue cohesivity may also be relevant for explanation of the MD. Figure 4D shows a voxel with tightly-packed tissue with collagen featuring an underestimated MD (blue border) and a voxel with loose tissue and few vessels with overestimated MD (purple border). This overestimation is more pronounced for the CD-based regression than for the CNN. An overview of residual maps of all samples can be found in the Supplementary material in Figures 4 and 5.

**Figure 4.**
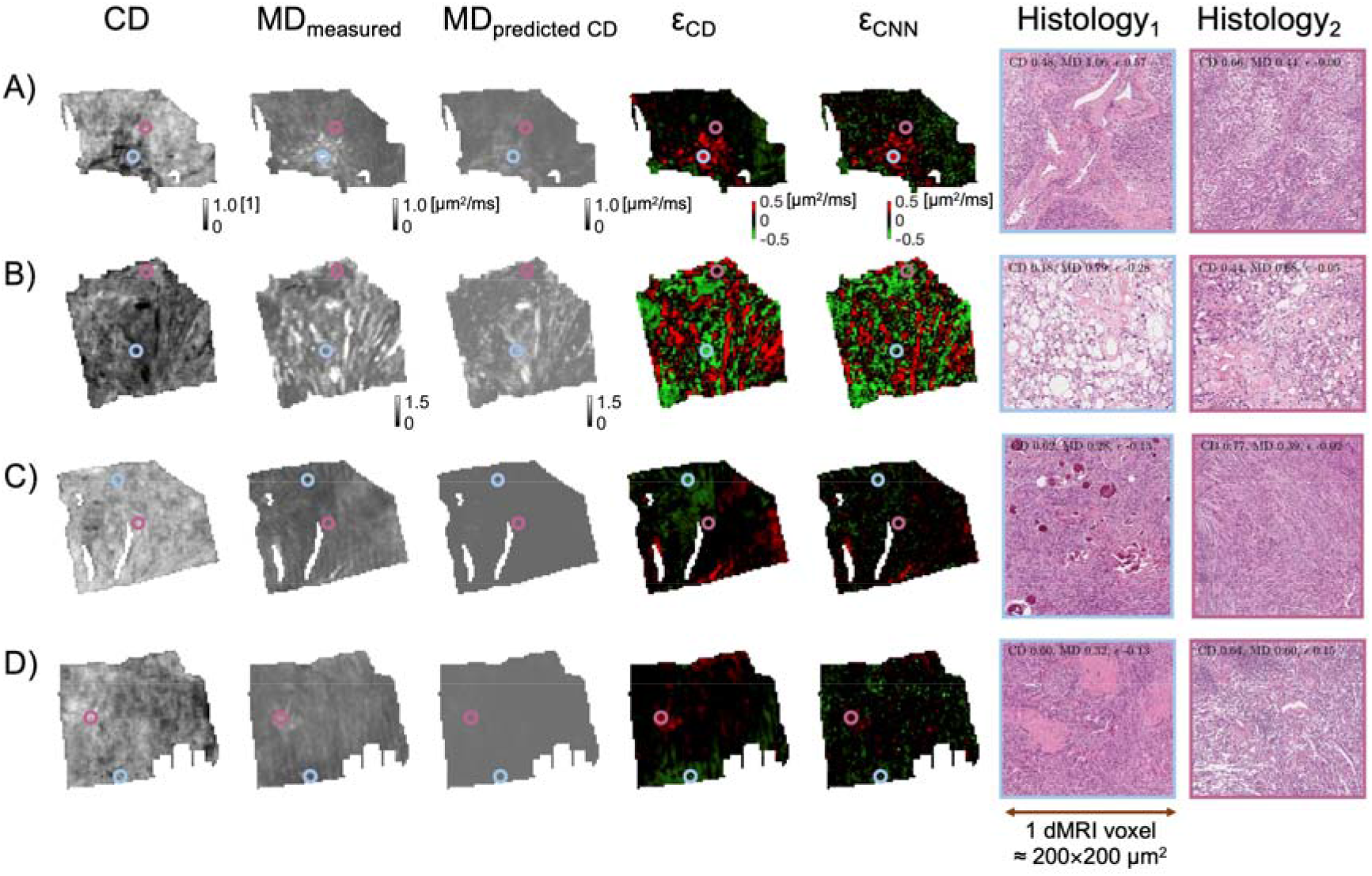
MD is influenced by histological features beyond CD. Columns show maps of the tumor sample and histology zoom-ins of a voxel with a feature associated with an MD poorly predicted by CD (Histology_1_) and a control voxel with an MD well-predicted by CD (Histology_2_). Two residual maps are shown – the first with the residual computed from MD predicted by CD (□_CD_) and the second with MD predicted by the CNN (□_CNN_). The color of the border of the zoom-ins matches the color of the circles, which indicate their origin in the sample. Panel A shows that a voxel with underestimated MD (red on the residual map; sample 7) contains tumor vasculature (blue marker), while the control voxel (purple marker) contains no large vessels but only tumor mass. Panel B shows a region with overestimated MD (green color on the residual map; sample 3) that can be linked to tightly packed microcysts (blue). The control voxel (purple) also features microcysts, but fewer. The residual around the microcysts (blue) appears to be dominant when CD is considered for its prediction (□_CD_) but not for CNN (□_CNN_). Panel C shows a region with overestimated MD (green on the residual map; sample 9) that could be attributed to psammoma bodies (blue). The control shows no psammoma bodies (purple). Panel D shows that MD can be linked to tissue cohesivity (sample 6). The overestimated voxel (green in the residual map) is associated with tightly-packed tissue with collagen (blue) whereas the underestimated region rather features loose tissue with few vessels (purple).

**Figure 5.**
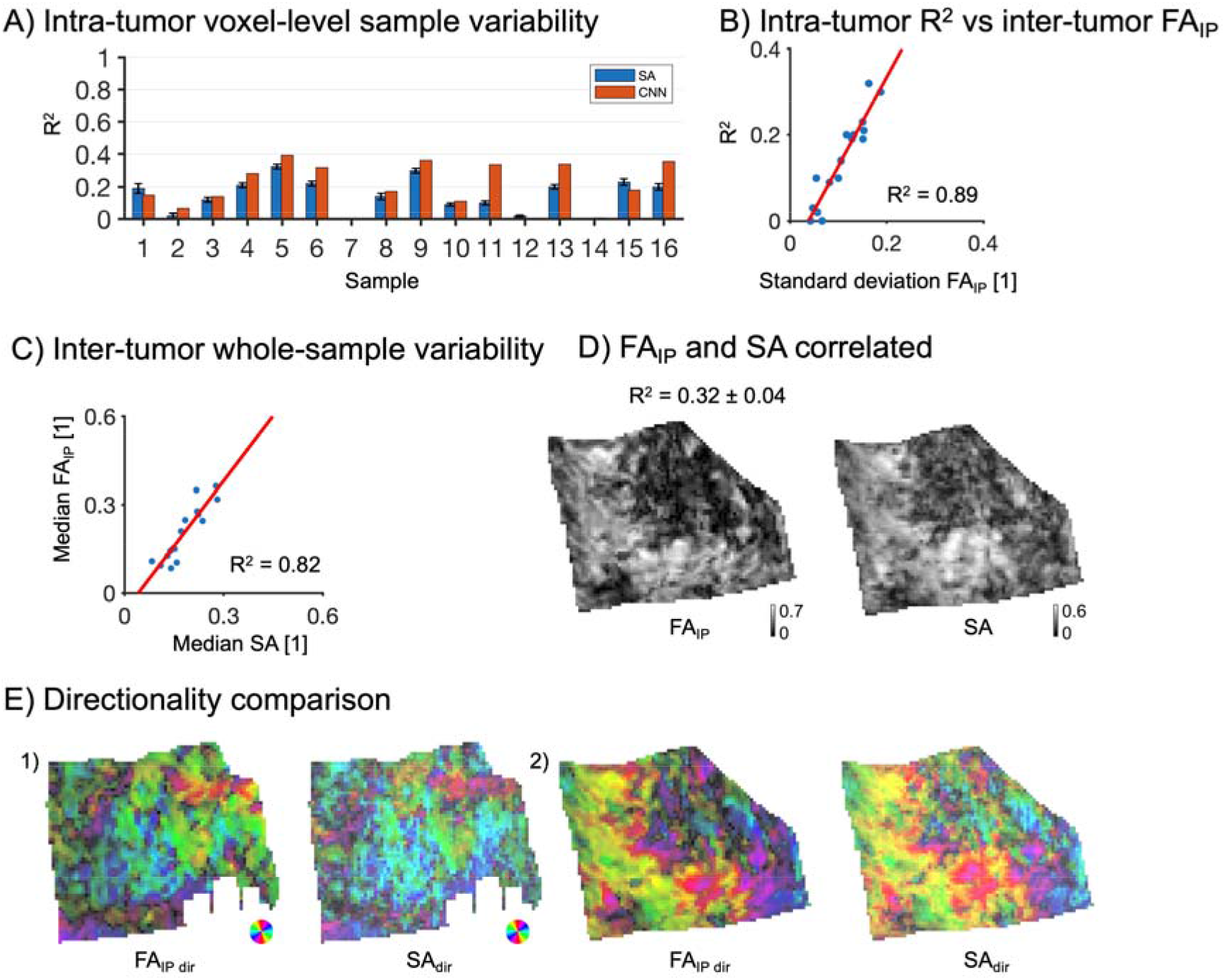
Association between FA and SA. Panel A shows the intra-tumor sample variability in FA_IP_ explained by SA (R^2^; blue bars correspond to median, black error bars show interquartile range) and by the convolutional neural network (CNN; red bars). Panel B shows R^2^ from the SA-based regression versus standard deviation of FA_IP_ values across the whole sample (each blue dot corresponds to a single sample; R^2^ = 0.89). Panel C shows FA_IP_ versus SA averaged across the whole sample and that SA explains FA_IP_ better than the intra-sample analysis (R^2^ = 0.82). Panel D shows a visual comparison of SA and FA_IP_ (sample 5). Panel E shows a comparison intensity is modulated by scaling the values by 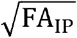 and 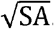.Visually, the between the directionality of SA and FA_IP_. Colors indicate directions, while the predicted and measured anisotropy and directionality of the samples are in strong agreement.

Just as for CD and MD, the SA explained FA_IP_ relatively poorly with R^2^ = 0.16 (0.06 – 0.20). However, here the per-sample R^2^ was strongly correlated with the standard deviation of FA_IP_ within samples (r = 0.94, p < 10^-5^, Pearson’s correlation coefficient), meaning that SA did predict FA_IP_ where there was sufficient feature-driven variation in FA_IP_ within the sample. The CNN displayed similar numbers as the SA-based regression, with R^2^ = 0.18 (0.09 – 0.34), however, there were a few samples where the performance of the CNN was much higher than that of the SA (3 samples: 11, 13, and 16). On the whole-tumor level, the association between FA_IP_ and SA was high with R^2^ = 0.82 (Figure 5C). From a visual perspective, the appearance of SA and FA_IP_ was similar, as illustrated for a sample with a high R^2^ of 0.32 (Figure 5D). The directionally encoded maps from dMRI and histology were also similar, as shown in two examples (Figure 5E). Corresponding maps for all tumors can be found in the Supplementary material.

To analyze the mechanism of the association between FA_IP_ and SA, histology images associated with MRI voxels with high or low FA_IP_ and SA were examined. Voxels with both high SA and high FA_IP_ featured elongated tissue structures oriented more or less along a single direction (Figure 6A), whereas voxels with both low SA and low FA_IP_ tended to feature high orientation dispersion where the mesoscopic organization appeared more disorganized (Figure 6B). Furthermore, some voxels featured high SA but low FA_IP_. Such voxels featured boundaries between tumor and vessels, transitions from tumor tissue to microcysts, or loose tissue with white transparent areas (Figure 6C), which yield high SA due to the strong contrast in the image but are in themselves not likely to have a strong effect on the diffusion. These voxels reflect a limitation in the use of SA as a proxy for tissue anisotropy.

**Figure 6.**
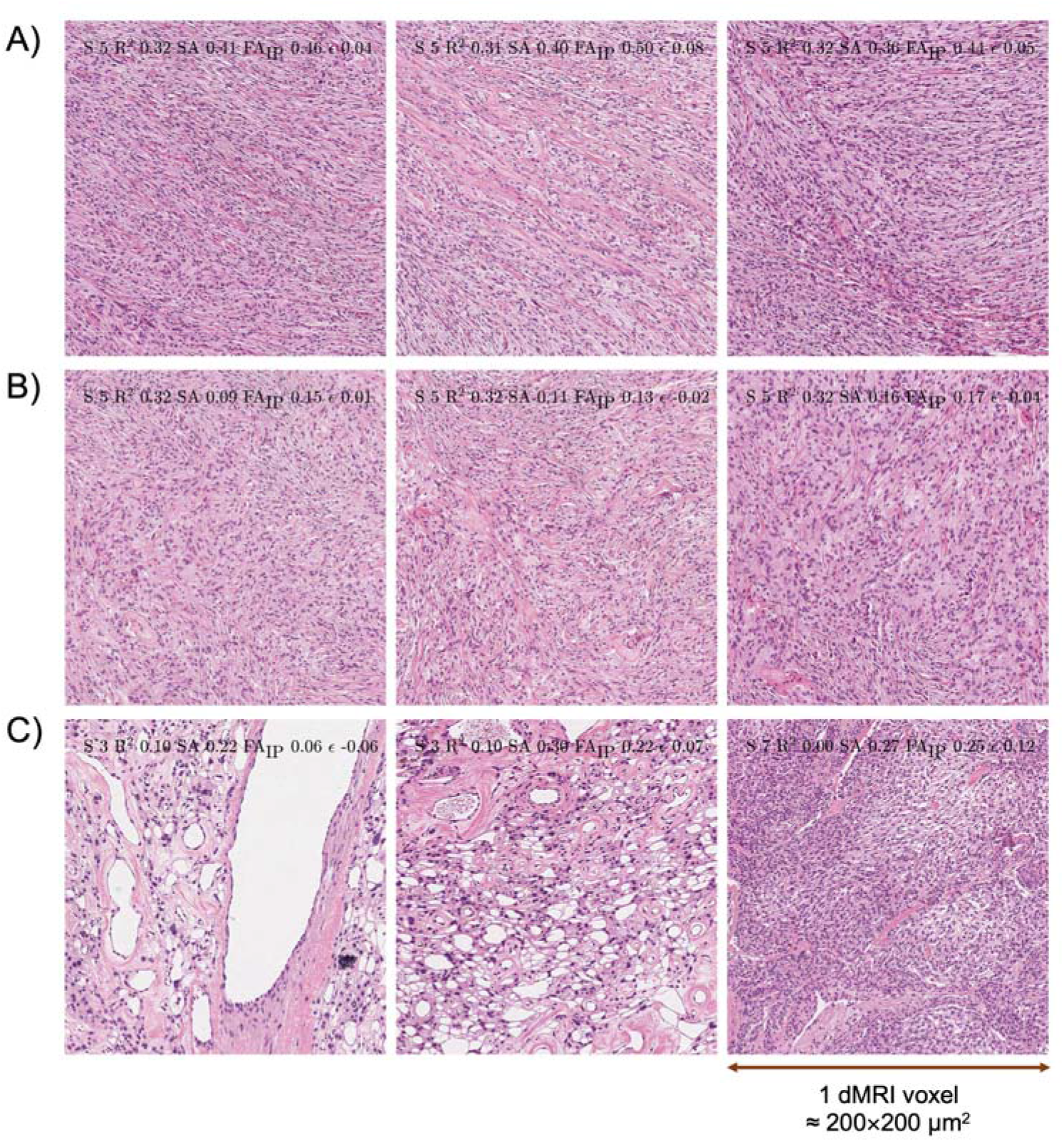
Histology that corresponds to MR voxels with high or low SA and FA_IP_. Panel A illustrates that tissue with elongated structures that are dominantly oriented along one direction yields high diffusion anisotropy (high SA and high FA_IP_). Panel B shows tissue structures oriented without any single preferential direction, which yield low diffusion anisotropy, and thus appear as isotropic tissue (low SA across the whole voxel and low FA_IP_). Panel C shows tissues with boundaries between tumor and vessels (left), transition from tumor tissue to microcysts (middle), or tissue looseness with white transparent areas (right). These yield high SA but the structures have little influence on the diffusion and thus yield low to intermediate FA_IP_. This illustrates a limitation of the structure tensor analysis technique.

## Discussion

We examined meningiomas ex vivo using both dMRI and histology in order to understand what microscopic and mesoscopic features of the tumor tissue that influences MD and FA from DTI. The analysis was applied to meningiomas of two different grades and six different types (Figure 1AB), which together displayed the highly heterogeneous microstructure typical for meningiomas (Wiemels et al., 2010). The data allowed us to test the common hypotheses that MD reflects cell density (CD) and that FA reflects tissue anisotropy as quantified by the structure anisotropy (SA). Results were in line with these hypotheses when analyzing data across tumors, however, the hypothesis did not hold within tumors. The cause of the discrepancy was different for MD and FA, however.

Regarding MD, the results indicate that CD alone is insufficient to explain the observed intra-tumor variability in MD. This is exemplified in Figure 3, where panels C and D show a case where CD is associated with MD whereas panels E and F show a case where it is not. In the third of the samples (6 out of 16), the CD was an exceedingly poor predictor of MD (R^2^ < 0.1). Across tumors, however, the MD did correlate negatively with CD (R^2^ = 0.58). The lack of an ability of CD to explain the intra-tumor variability in MD could be that the factor that determines MD is the intracellular volume fraction (ICVF) rather than cell density as defined as the number of cells per volume unit (example of histology shown in Figure 3F) (Szafer et al., 1995;Chenevert et al., 2000;Nilsson et al., 2018a;Novikov et al., 2019). This is because the MD is determined by the volume-weighted average of diffusivities in the intra- and extracellular spaces. For cells smaller than approximately 10–15 µm, the intracellular MD is close to zero (Szafer et al., 1995). Therefore the MD on a voxel-level is given by the product between the extracellular volume fraction (given as 1– ICVF) and the extracellular MD. Voxels with intermediate and high CD could have similar ICVF if their cell sizes were different (e.g. Figure 3F), and thus similar MD according to this conceptual model. Voxels with very low CD, however, tend to feature loose and necrotic tissue, which leads to lower ICVF and thus higher MD. The comparison across tumors on the whole-sample level supports this idea since on this level the association between CD and MD is driven by the tumors with low to intermediate CD while for intermediate to high CD there is no discernable association (Figure 3B). Another reason why CD generally failed to explain MD could be that for many samples the measured MD showed little to no variations across the sample, which means that there was no variation to explain (coefficient of variation in MD was below 0.2 for 10 out of 16 samples; see Figures 1 and 2 in the Supplementary material). This means that any microscopic feature would yield low R^2^. However, the CD showed considerable variation in many such cases (coefficient of variation in CD was below 0.2 for only 5 out of 16 samples; see also Supplementary material). This means different CD gives highly similar MD in many samples, which emphasizes the argument raised above. Furthermore, the CNN explained more variance in MD than the CD, which suggests that features apart from CD contribute to the variation in MD. Examples of such features, identified by inspection of residual maps, include tumor vasculature, psammoma bodies, microcysts, and tissue cohesivity (Figure 4). We hypothesize that these are relevant features for MD because MD is poorly explained by CD alone, as well as because MD prediction is less biased when the more general and flexible CNN approach is used. This is also in agreement with other studies arguing that features of the mesoscopic stromal architecture influence MD more than CD (Squillaci et al., 2004;Yoshikawa et al., 2008). Furthermore, stromal collagen content (Egnell et al., 2020) or the presence of necrosis may also influence MD (Patterson et al., 2008). Modelling work shows that MD can also be influenced by features of the cells such as their size (Szafer et al., 1995), nuclear size (Xu et al., 2009), or membrane permeability (Colvin et al., 2011). This work hypothesizes that other microstructural features are of importance (Figure 4) but quantifying their effects will be the subject of future work.

The results concerning anisotropy were seemingly similar to those concerning cell density, but subtle differences offer a different interpretation. Similar to MD and CD, FA_IP_ was better explained by SA on the inter-tumor level than on the intra-tumor level (Figure 5AC). Samples with fewer variations of FA_IP_ had markedly lower R^2^ values (Figure 5B). This is because the relative importance of noise is higher for samples with low FA_IP_ variability. On the other hand, samples with high FA_IP_ variability have more true variation that needs to be explained compared to noise. Importantly, samples with a uniform and low FA also showed uniform values and low values of the SA. Note the difference from the case of MD and CD, where a uniform MD was found even in samples with a non-uniform CD. Furthermore, high SA and high FA_IP_ were associated with the presence of anisotropic tissue structures, and low SA and low FA_IP_ with either isotropic tissue structures or anisotropic tissue structures with high orientation dispersion (Figure 6AB). This is aligned with prior research (Pierpaoli et al., 1996;Szczepankiewicz et al., 2016), because FA corresponds to the voxel-level average diffusion anisotropy, which is high only in the presence of aligned and elongated microscopic structures and low if either microscopic diffusion anisotropy is low or orientation dispersion is high or both (Szczepankiewicz et al., 2016).

Gaining detailed knowledge of tumor microstructure non-invasively by diffusion MRI is a desirable goal, however, our results show that MD and FA are affected by a multitude of different microstructure features and thus lack specific interpretations. To enable the separation of the many features that affect MD of FA, we need to use diffusion protocols that encode more information than standard DTI protocols (Nilsson et al., 2018a). For example, time-dependent diffusion (Stepišnik, 1993) could potentially be used to distinguish microcysts from CD because microcysts are circumscribed by an endothelial layer, and their sizes are on average larger than cells. Strong effects of diffusion time could thus indicate the presence of microcysts. Tensor-valued diffusion encoding may also be used to encode for microscopic anisotropy that is independent of tissue orientation dispersion (Szczepankiewicz et al., 2015;Szczepankiewicz et al., 2016;Westin et al., 2016).

In this study, we identified six potential limitations of the present work. First, the ability to use histology to predict dMRI parameters depends on the accuracy of image coregistration. Herein lies an intrinsic limitation as the MRI voxels were 200 µm thick, whereas the histology sections were only 5 µm thick. The sections were also somewhat deformed during preparation. The influence of the latter limitation was limited by performing both linear and non-linear registration between the histology images and the dMRI maps. Nonetheless, some of the large residuals seen in highly heterogeneous samples (e.g. Figure 3C) could be due to a spatial mismatch in the through-slice direction between the two modalities. However, such a mismatch is unlikely to have affected the residual maps in Figure 4A, 4C and 4D, where highly localized and sample-specific features were clearly related to the residuals.

Furthermore, potential registration errors are unlikely to have affected the low R^2^ in samples with uniform MD or FA_IP_, as these simply lacked variance to be explained. A second limitation is that features beyond CD that affects MD were identified only qualitatively. Further work is needed to enable quantification of those features in order to quantify the strength of their association with MD. A third limitation is that the meningioma classification was based on Louis et al. (2016) although a newer classification was proposed after the study was closed (Louis et al., 2021). However, the classification was not used in the analysis. A fourth limitation is that the analysis used a second order polynomial to relate histological image features with measured dMRI parameters. Such a polynomial lacks a biophysical foundation but explained the data reasonably well and served our goal to test for an association between CD and MD or SA and FA_IP_. Future work could possibly use biophysical modelling to better relate histology to MRI. A fifth limitation is that the training of the CNN only adjusted a limited number of parameters in the final prediction layers. Training a convolutional neural network without using pre-trained network could yield better performance, but prior work reported that fine tuning of networks pretrained on large sets of images yields better performance for a histology classification task (Vesal et al., 2018). Finally, the results were obtained ex-vivo which may not fully generalize to the in-vivo situation.

## Conclusion

The association between MD and cell density was present only when comparing across tumors. On the mesoscopic level within tumors, the MD in meningiomas was not determined by the cell density, as several samples with highly variable cell density but uniform MD were found. We argue that, on the mesoscopic level, the MD may be influenced by the intracellular volume fraction rather than the cellularity. The MD is also influenced by other features such as the presence of large vessels, microcysts, psammoma bodies, and the looseness of the tissue. Furthermore, FA was linked to the tissue structure anisotropy and we found support that it is elevated in the presence of elongated and aligned cell structures in line with previous knowledge.

## Abbreviations

CD: Cell density or cellularity
CNN: Convolutional neuronal network
DEC: Directionally encoded color maps
dMRI: Diffusion magnetic resonance imaging
DTI: Diffusion tensor imaging
EPI: Echo-planar imaging
FA: Fractional anisotropy
FA_IP_: In-plane fractional anisotropy
H&E: Hematoxylin & eosin
ICVF: Intracellular volume fraction
MD: Mean diffusivity
SA: Structure anisotropy
WHO: World Health Organization

## Acknowledgment

Michael Gottschalk, René In ‘T Zandt and Lund University Bioimaging Center (LBIC), Lund University are gratefully acknowledged for providing experimental resources.

## Credit author statement

**Jan Brabec**: Conceptualization, Methodology, Formal analysis, Investigation, Formal analysis, Visualization, Data Curation, Writing - Original Draft, Writing - Review & Editing **Magda Friedjungová**: Software, Formal analysis, Data Curation, Conceptualization, Writing - Review & Editing **Daniel Vašata**: Software, Formal analysis, Data Curation, Conceptualization, Writing - Review & Editing **Elisabet Englund**: Resources, Investigation, Project administration, Writing - Review & Editing **Johan Bengzon**: Resources, Funding acquisition, Project administration, Writing - Review & Editing **Linda Knutsson**: Resources, Funding acquisition, Writing - Review & Editing **Filip Szczepankiewicz**: Conceptualization, Methodology, Software, Supervision, Writing - Original Draft, Writing - Review & Editing **Pia C Sundgren**: Funding acquisition, Project administration, Writing - Review & Editing **Markus Nilsson**: Conceptualization, Funding acquisition, Visualization, Methodology, Software, Project administration, Supervision, Writing - Review & Editing, Visualization.

## Tables

**Table 3.**
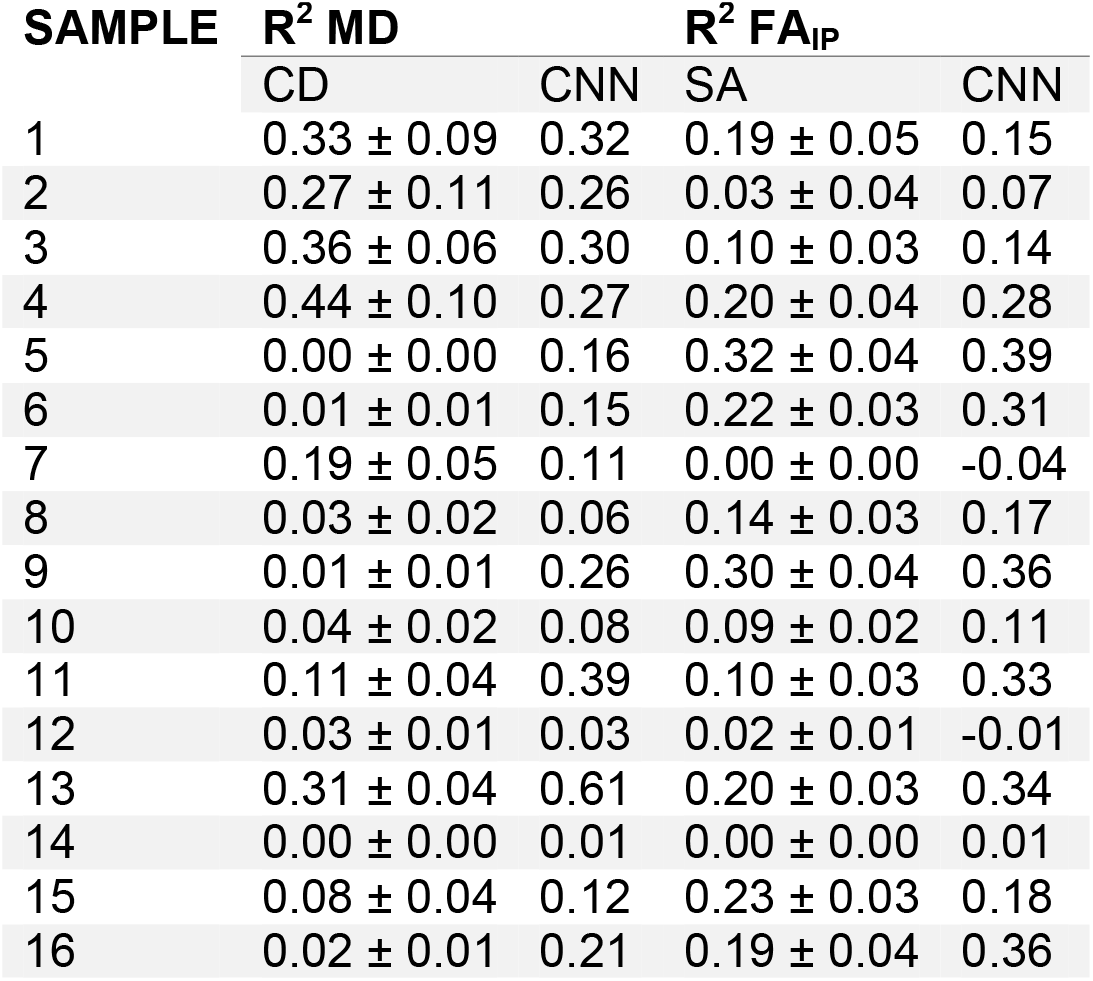
Coefficient of determination values from the test sets between measured and predicted values. R^2^ values are in the format median ± interquartile

