## Supplementary material for "Meningioma microstructure assessed by diffusion MRI: an investigation of the source of mean diffusivity and fractional anisotropy by quantitative histology"

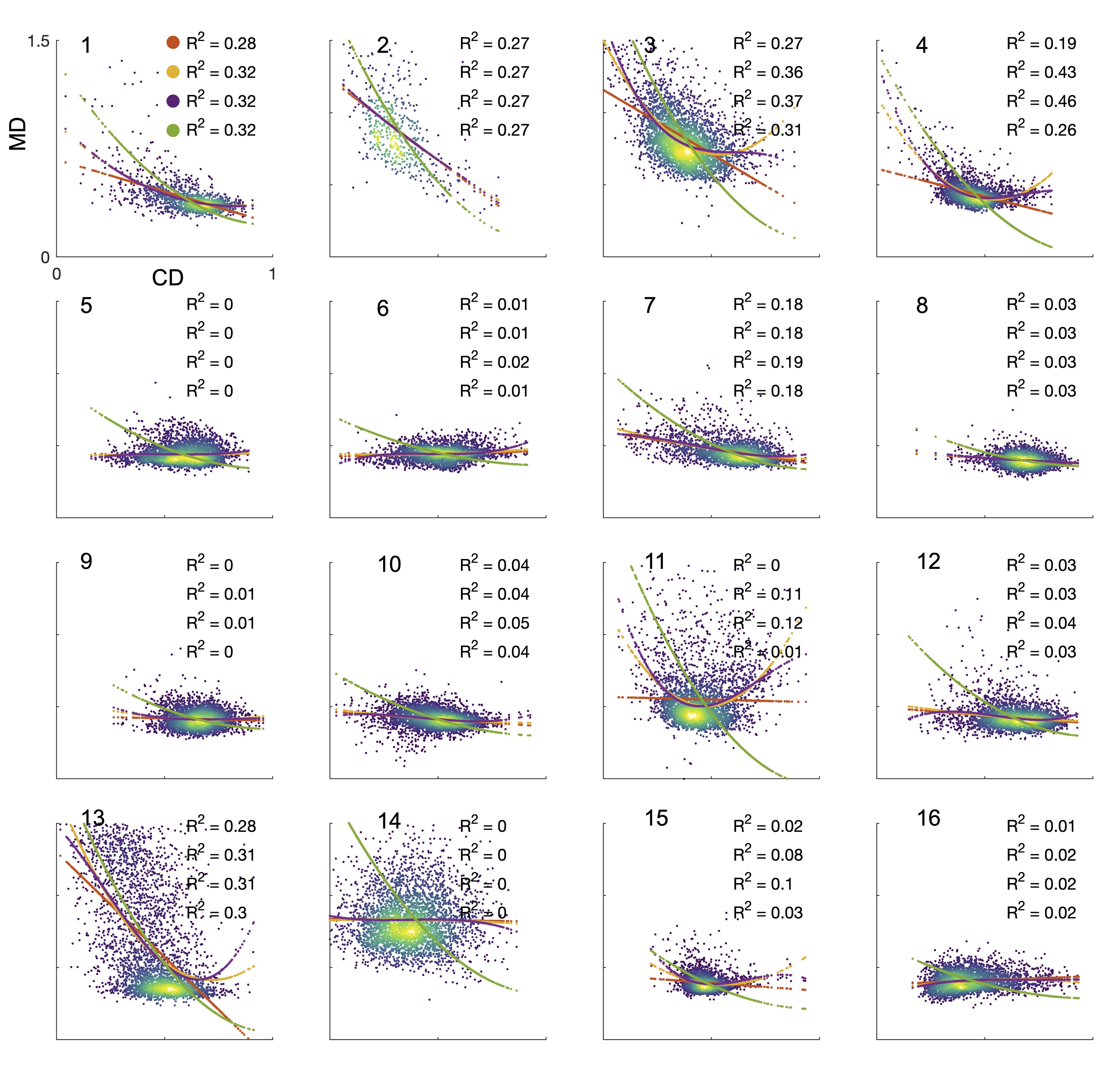

**Supplementary Figure 1. Scatter plots and fits of CD vs MD of all samples.** Red color represents a first-degree polynomial, orange color represents a second-degree polynomial and purple color represents a third-degree polynomial. Green color represents a second-degree polynomial constrained to maximal values at minimal CD and monotonically decreasing. Across all samples, we settled upon the second-degree polynomial for further analysis. Sample number in the part of the scatter plot and R2 on the right part of the scatter plot (top row linear, second second-degree, third-degree and last row second-degree constrained polynomial).

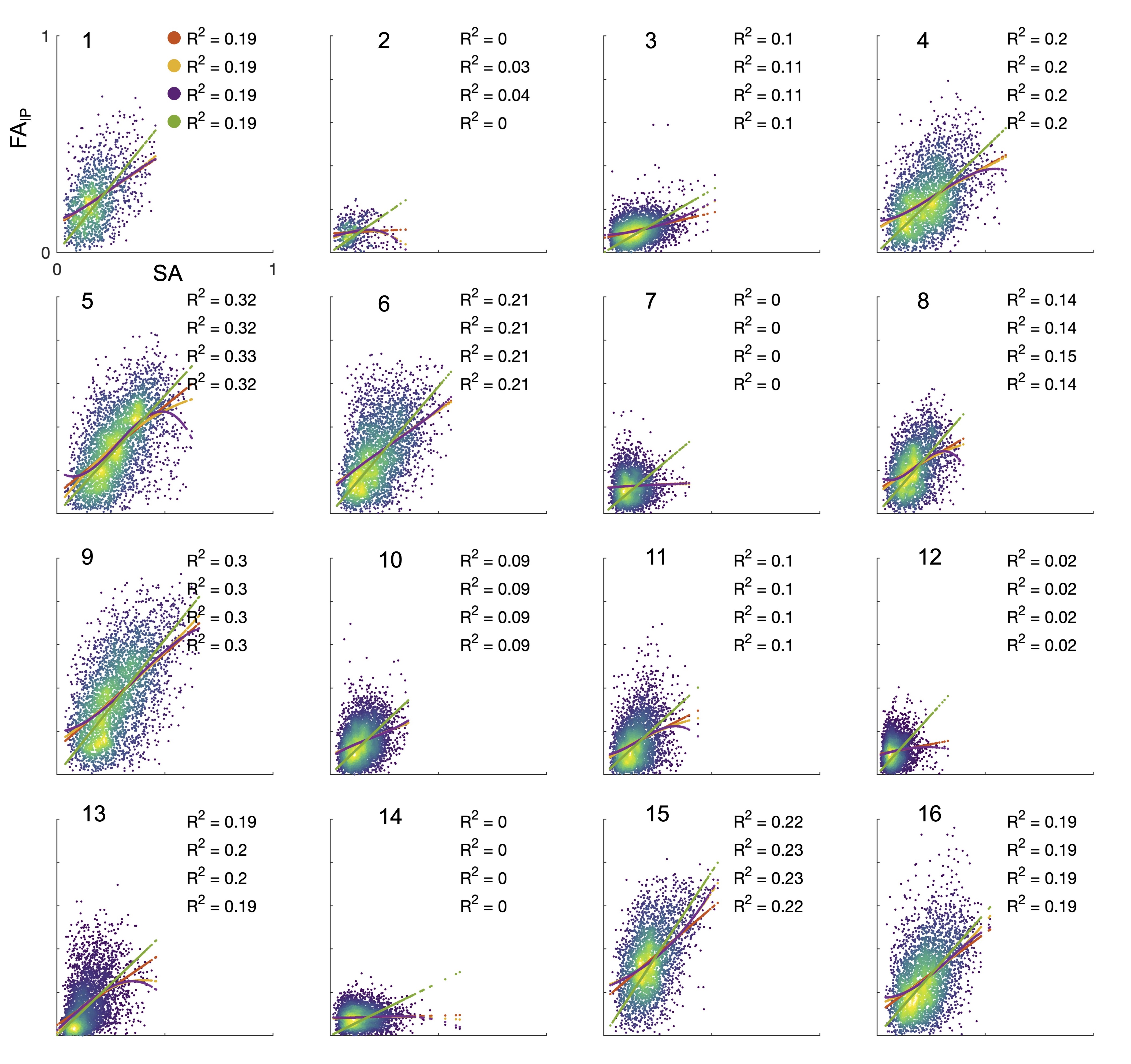

**Supplementary Figure 2. Scatter plots and fits of SA vs FAIP in all samples.** Red color represents a first-degree polynomial, orange represents a second-degree polynomial, purple represents a third-degree polynomial and green represents a second-degree polynomial constrained to the origin (0,0). Across all samples, we settled upon the second-degree polynomial for further analysis. Sample number in the part of the scatter plot and R2 on the right part of the scatter plot (top row linear, second second-degree, third-degree and last row first-degree constrained polynomial).

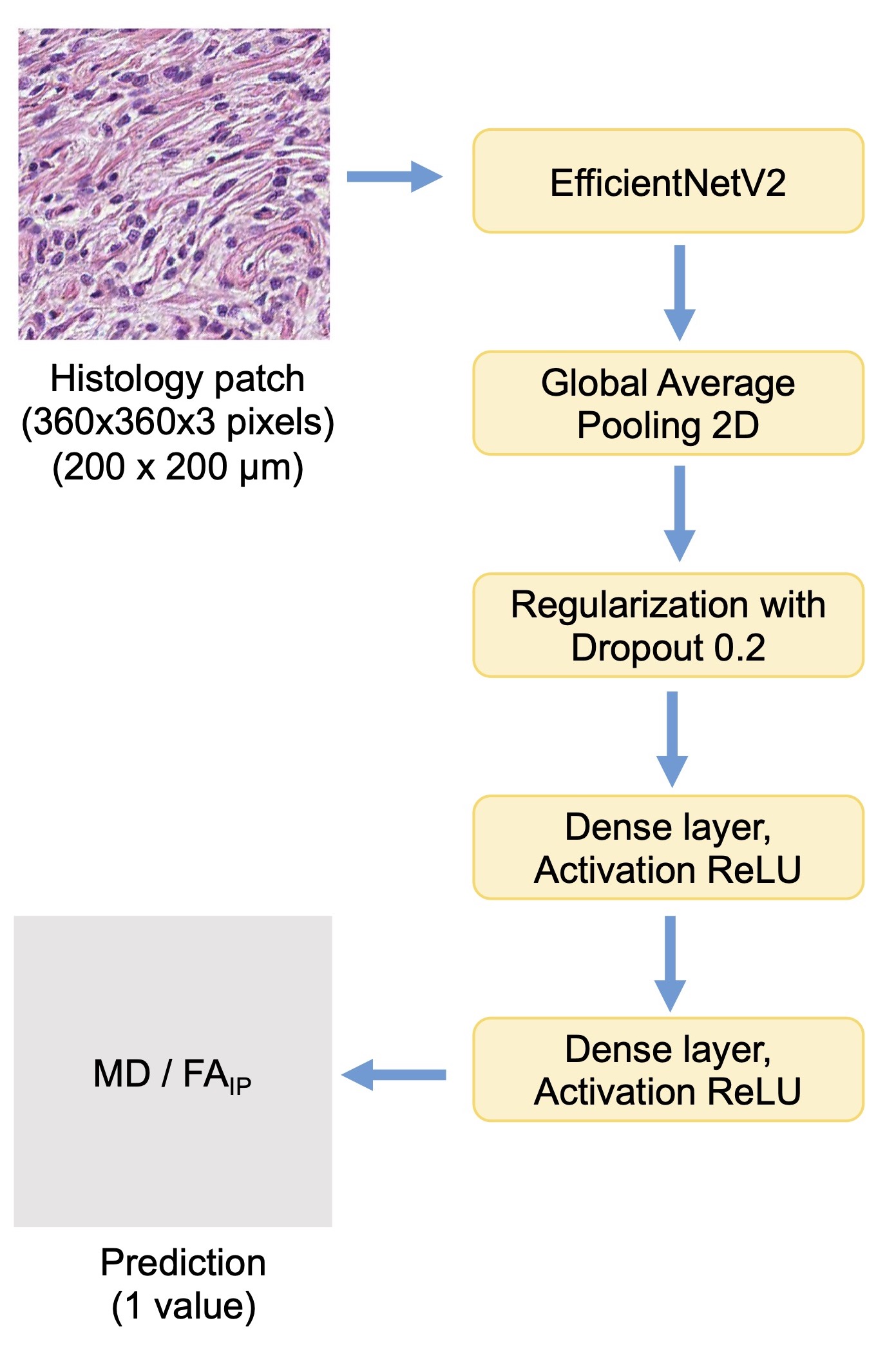

**Supplementary Figure 3. An overview of the convolutional neural network (CNN) architecture.**

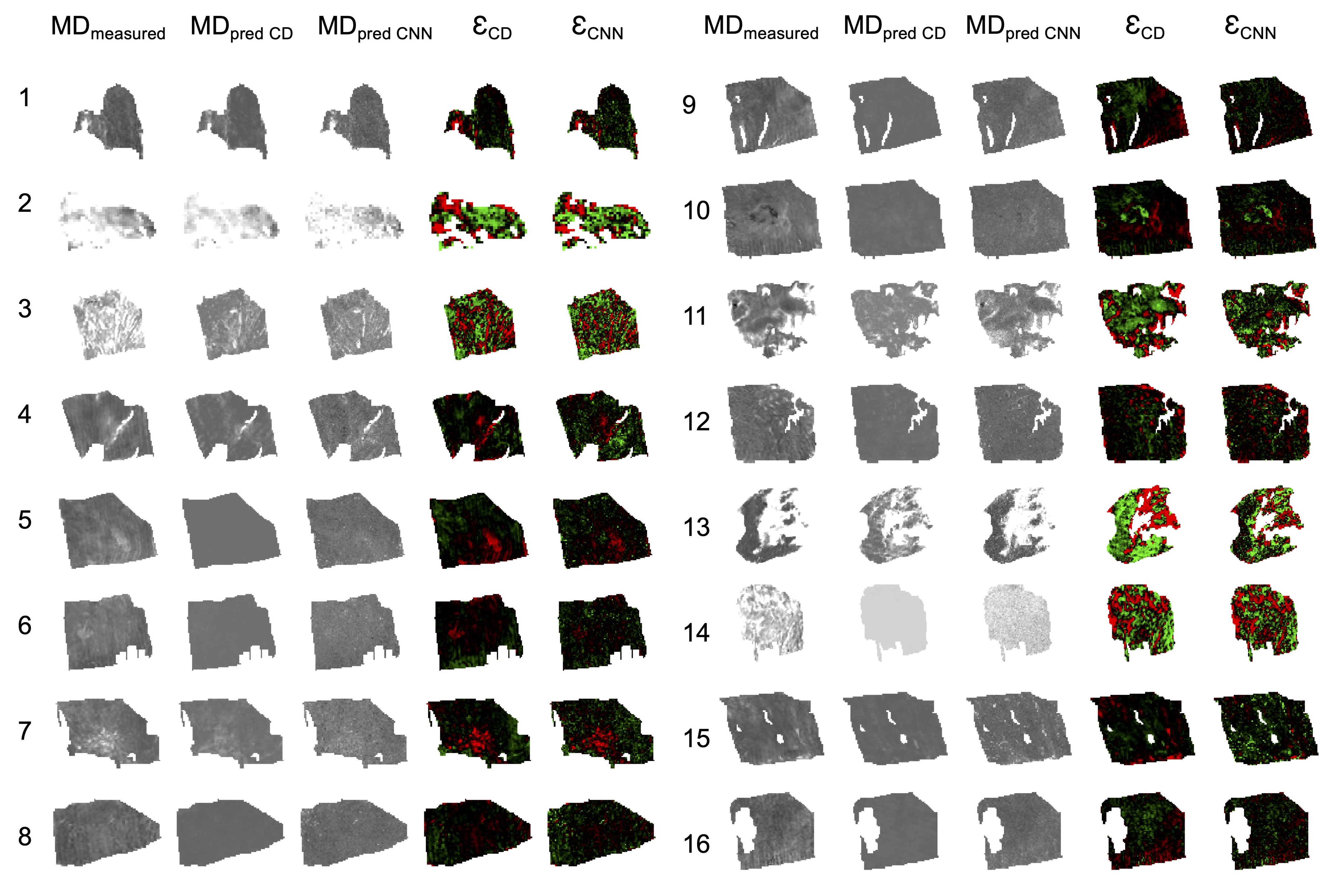

**Supplementary Figure 4. Predicted MD by cell density (CD) and convolutional neural network (CNN) together with residual maps.** The figure shows an overview of all samples with measured MD and MD predicted by CD (MD_pred CD_) or CNN (MD_pred CNN_) as well as their residual maps (measured – predicted MD) of CD (Ɛ_CD_) and CNN (Ɛ_CNN_) predictions.

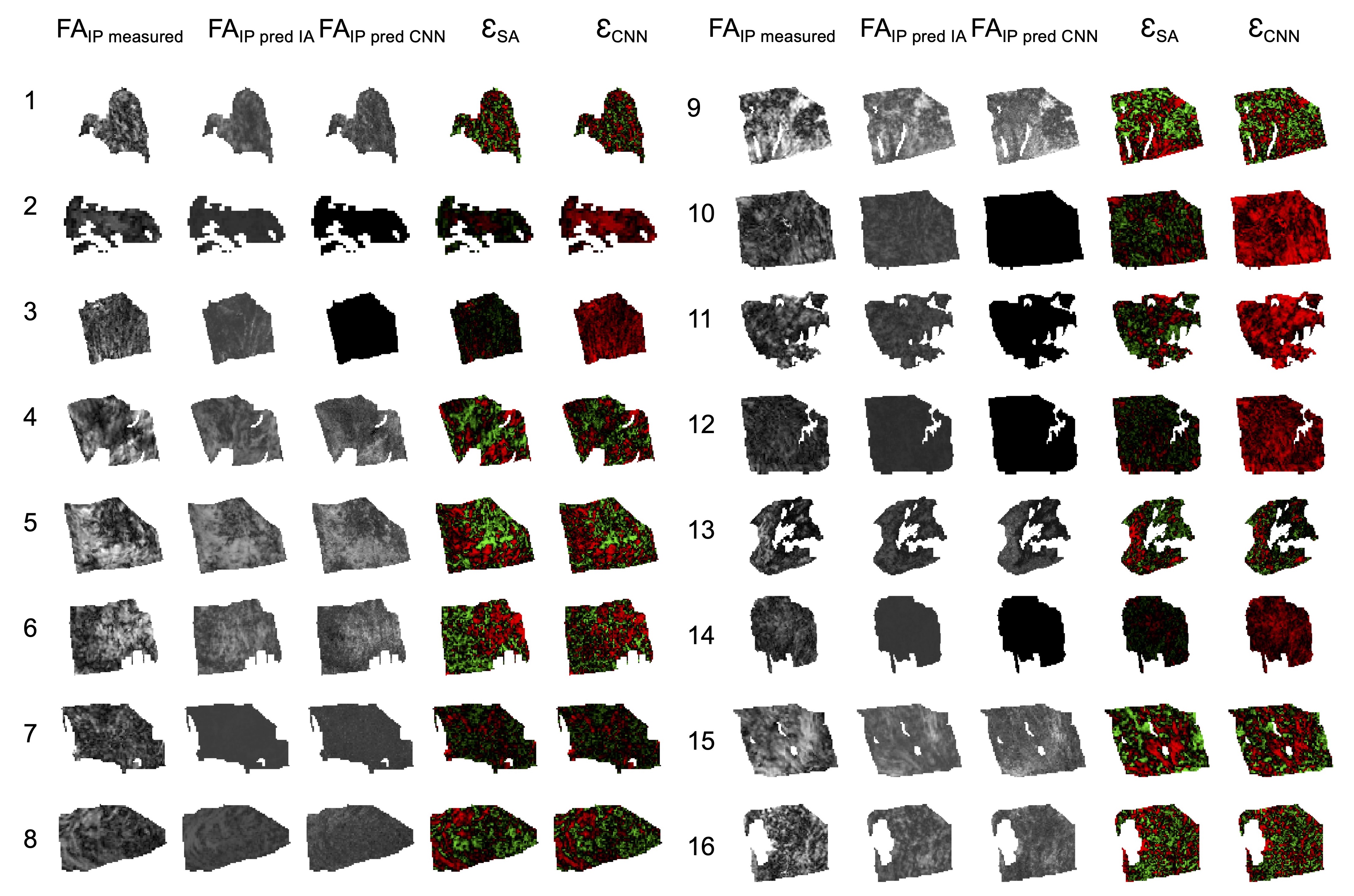

**Supplementary Figure 5. Predicted FA_IP_ by structure anisotropy (SA) and convolutional neural network (CNN) together with residual maps.** The figure shows an overview of all samples with measured FA_IP_ and FA_IP_ predicted by CD (FA_IP_ pred CD) or CNN (FA_IP_ pred CNN) as well as their residual maps (measured – predicted FA_IP_) of SA (Ɛ_SA_) and CNN (Ɛ_CNN_) predictions.

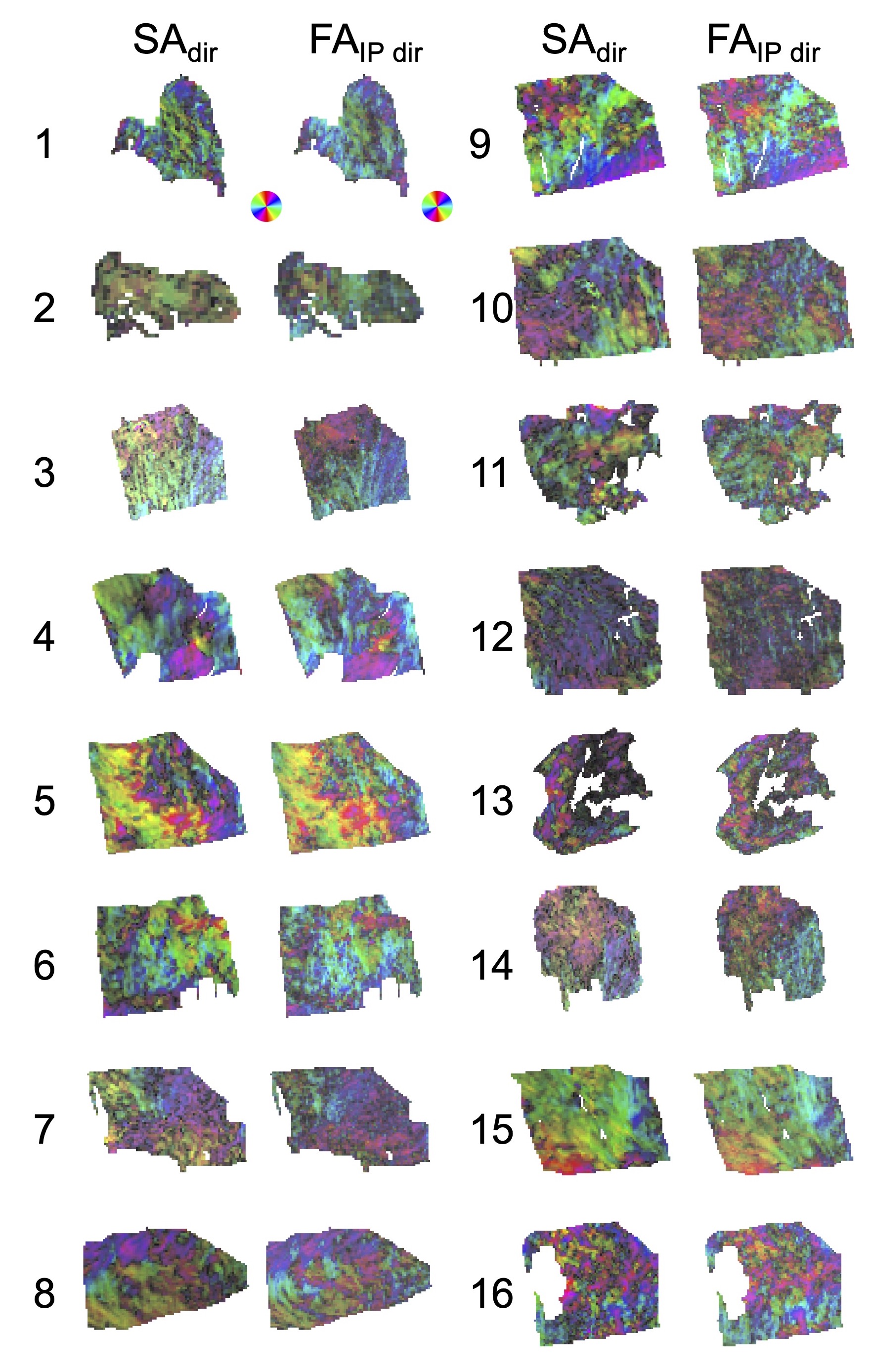

**Supplementary Figure 6. Comparison between directionality of SA and FA_IP_.** The directions are color-coded and the intensity of FA_IP_ and SA is suppressed by scaling the values by $\sqrt{FA_{\mathrm{IP}}}$ and $\sqrt{\mathrm{SA}}$.
